# Divide and Conquer approach for Genome Classification based on subclass characterization

**DOI:** 10.1101/003475

**Authors:** S.S. Patil, M.N. Murty, U.B. Angadi

## Abstract

Classification of large grass genome sequences has major challenges in functional genomes. The presence of motifs in grass genome chains can make the prediction of the functional behavior of grass genome possible. The correlation between grass genome properties and their motifs is not always obvious, since more than one motif may exist within a genome chain. Due to the complexity of this association most pattern classification algorithms are either vain or time consuming. Attempted to a reduction of high dimensional data that utilizes DAC technique is presented. Data are disjoining into equal multiple sets while preserving the original data distribution in each set. Then, multiple modules are created by using the data sets as independent training sets and classified into respective modules. Finally, the modules are combined to produce the final classification rules, containing all the previously extracted information. The methodology is tested using various grass genome data sets. Results indicate that the time efficiency of our algorithm is improved compared to other known data mining algorithms.

## 1 Introduction

The development of elaborated and specialized bioinformatics computational tools has led to revolutionary changes in the analysis of genome sequences. Visualizing, manipulating and predicting molecular structure and function, separating DNA sequence according to grass genome coding regions, classifying grass genome, DNA and RNA molecules or detecting weak similarities have come to rely vitally on computational methods [3]. The continuous increase in size of biological genomic databases requires new and computation-sensitive data mining approaches, and also presents unique opportunities for new fields of inquiry; among these lies one of bioinformatics more ambitious goals, the prediction of the functional behavior of grass genome. Genome are large molecules composed of one or more chains of nucleic acids in a specific order, which is determined by the base sequence of the nucleotides in the gene coding of the grass genome.

A DAC technique divide fragments of grass genome sequences large data sets into k/n sub sets. Common behavioral characteristics and strong structural similarities enable classification of genomes into genome families. The lack of cost and time-efficient experimental methods has given rise to computational approaches for grass genome properties identification. Moreover, the overlapping of grass genome families is almost at all times, the classification of a grass genome into multiple families with different similarity levels makes the procedure even harder, due to its ascending complexity. Despite the preceding difficulties, grass genome functionality prediction could hardly be achieved not only for *motifs*, short nucleic acid sequences of specific order, which appear in grass genome chains and play a decisive role in grass genome behavior. Although a straightforward mapping between motifs and grass genome properties is hard to achieve due to the presence of multiple motifs in each grass genome chain, they can facilitate prediction of grass genome functionality, if the latter is considered to be derived by the combining effect of many, either conflicting or consistent motifs. Two different types of motifs can be distinguished into *patterns* and *profiles*. The first is the simplest form of a motif and can be represented by an expression. More complex than a pattern, the profile constitutes a multiple sequence alignment, symbolized by alignment matrices. The question of occurrence of a particular motif in a grass genome chain is obtained when a pattern motifs counter is incremented. The overall procedure of motif identification and detection in a genome sequence can be carried out in two different ways: *unsupervised* or *supervised*. The former can be accomplished by a large variety of machine learning algorithms, whereas the latter requires prior information, such as expert opinion or experiment conclusions. Germ plasma, Entrez, Gramene. Related literature features many data mining algorithms that utilize the presence of motifs in genome sequences to perform genome classification, originating from the field of pattern recognition [7], as well as that of artificial intelligence [6]. They include many different techniques, such as decision trees [14], statistical models, neural networks [5] and Grid Classification [11]. This paper presents DAC (*Genome Classification*), a novel methodology that aims to benefit from the use of DAC technology in data mining applications. The combination of the two leading technologies can help overcome the computational difficulties often encountered in genome classification problems. The Genome DAC Class methodology follows a “divide and conquer” approach comprising three steps: First, Grass Genome data from an expert-based database are divided into multiple disjoint sets, each one preserving the original data distribution. Next, the new sets are used as training sets, and multiple modules are derived by clustering the set of sequences of their motifs with hamming distances means of standard data mining algorithms [15]. Finally, the modules are combined to produce the final classification rules, which can be used to classify a given instance and evaluate the methodology.

## 2. Methodology Outline

The main goal of the DAC methodology is to utilize existing data mining algorithms in a parallel-enabled environment, such as the DAC, in order to create a Grass Genomes classification module. Divide-and-conquer technique is a powerful problem solving technique that is the basis for many effective sequential algorithms. We analyze the extent to which divide-and-conquer yields effective and efficient parallel algorithms. We identify number of equivalence classes of divide-and-conquer algorithms and determine which classes are good candidates for parallelization and the architectures for which they are best suited. None of the classes provide optimal speedup when the maximum possible numbers of processors are used, and four of them yield no more than constant speedup. However, eleven classes do provide optimal speedup under limited parallelism, and three of these classes have polylogarithmic runtime with a polynomial number of processors and they accelerate within a polylogarithmic factor of optimal under maximum parallelism. The communication cost incurred during parallelization is found to have a significant impact on the performance of a parallel divide-and-conquer algorithm. This factor alone can mean the difference between expedite that is within a poly logarithmic factor of optimal and no speedup at all.

### 2.1. Data Splitting

Divide the grass genome sequences dataset into number of sub datasets using splitting technique. The first step of the DAC methodology, after the creation of the original single dataset, is to divide it into multiple datasets. The process is performed by the algorithm Splitter presented. As input, the algorithm requires datasets with the data to be divided. It then reorders the data so as to cluster entries with the same grass genome class. Finally, instead of simply chopping the sequences data into equal parts, datasets employs a method similar to Round-Robin to equally distribute the members of each class to all the dataset, thus preserving the initial class distribution. Experimentally, it is found that the method is robust in the class allocation, both for different number of splits and for varying number of classes involved. The splitting of the original data being complete, the next step in the DAC methodology is creating of the individual knowledge modules. Each data set is used to train and test a separate classifier. Since each dataset contains a disjoint subset of the original data, they can be processed in parallel for time efficiency. Any classification algorithm that can be applied on nominal values can be used in this phase. Using the Leader is an incremental algorithm in which L leaders (Lds) each representing a cluster is generated using a suitable threshold value. As an extension of leader algorithm, we have implemented Leaders algorithm. In this method, after finding L leaders using the leader algorithm main leaders are generated within each cluster represented by a leader, choosing a suitable sub threshold value. Thus, Leaders creates L clusters with L leaders in the i^th^ cluster as shown in Fig. 1. Sub leaders are the representatives of the sub clusters and they in turn help in classifying the given new/test pattern more accurately.

**Fig-1.**
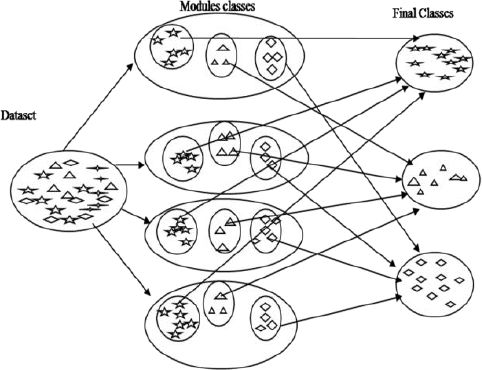
Divide and conquer methodology, Clustering and Classification

This algorithm can be used to generate a hierarchical structure as shown in Fig.1 and this procedure may be extended to more than two levels. An each k level hierarchical structure can be generated in only k database scans and is computationally less expensive compared to other hierarchical clustering algorithms. Number of database scans is < k since the number of training patterns to be scanned during clustering decreases as k increases and also we can efficiently manage the memory. If a representative from each subgroup is chosen then naturally classification accuracy (CA) would be improved. Hamming distance is used for characterizing dissimilarity between two patterns in case of grass genome data set. Hamming distance between two, d dimensional patterns :x and y is given by, threshold values can be initially chosen depending on the maximum and the minimum hamming distance values between the objects of a class in case of supervised learning. For unsupervised clustering technique, threshold value should be chosen properly depending on the number of clusters to be generated. If the threshold value is too small then a large number of clusters are generated and if the threshold value is too large then very few clusters are generated. Prototypes (representatives of the clusters and sub clusters) are generated using the training data set. During classification/testing phase, for every test pattern of the testing data set, the nearest leader is found first and then the nearest sub leader in that cluster is determined. Then the test pattern is classified based on the nearest of these two. For different threshold and sub threshold values, experiments (both training and testing) are conducted and the results are evaluated. To evaluate the clustering quality (quality of the prototypes selected) labeled patterns are considered. During training phase they are treated as unlabelled patterns and prototypes are selected. The quality of the prototypes is evaluated using the CA obtained for the testing data set [1]. Class-Subclass algorithm require two database scans (k=2) and its time complexity is O(*ndk*) and is computationally less expensive compared to most of the other partitional and hierarchical clustering algorithms as shown in Table 2. Space complexity and the sum of the prototypes is less than the total number of patterns n. The space requirement will be reduced as only these representatives are to be stored in the main memory during the testing phase. Even if more number of prototypes is generated, classification time is less as only part of the hierarchical structure is searched during the testing phase

### 2.2. The Classifier Training process on DAC Computing

The splitting of the original data being complete the next step in the DAC methodology is creating of the individual knowledge modules. Since each dataset contains a disjoint subset of the original data they can be processed in parallel for time efficiency. Any classification algorithm that can be applied on nominal values can be used in this phase. In order to facilitate an efficient way to train the multiple modules simultaneously, the training phase makes use of the DAC resources. The grid genome sequences dataset concept is created in order to provide a distributed computing infrastructure for advanced science and engineering [9]. It aims to facilitate the flexible, secure, coordinated and controlled resources sharing among computers, to distribute computing power, data storage and specialized equipment use among the computing nodes scattered all over the world and assist the virtual organization, dynamic collection of individuals, groups. The grid offers the resources needed to run multiple training processes, thus reducing the total time cost of the classification procedure. The grid infrastructure is then responsible for assigning the suitable resources according to the description in the DAC grass genome dataset and queue the appropriate Grid node after the successful execution of the training process, the resulting output files returned. This process is repeated for the multiple training processes, each one of them assigned to Grid node for execution. At this point it must be noted that, contrary to the other parallel classification techniques, the DAC methodology is independent of the actual data mining algorithm. Any classification algorithm available in literature can be utilized in the classifier training process, thus providing a degree of freedom to the methodology. Also, due to the fact that the training dataset were equivalent regarding the actual data representation, the final knowledge modules are also of equivalent accuracy.

### 2.3. Merging the Classified datasets

The efficient combination of the multiple knowledge modules, were extracted in the previous section. The modules are combined by a process, where each one is represented by a distribution vector. The performance of the total classifier for a single instance is derived by the average of all the distribution vectors. The overall efficiency of the module is calculated by testing it on the original dataset. All Ci classes classified from sub clusters are merged to form a classified single data set. Repeating the procedure of sub leaders algorithm to make the clusters, and using nearest neighbor classifier with hamming distance similarity measure the sub classes are obtained. The algorithm used is presented in more detail in algorithm. Results found through extensive experimentation have shown that the accuracy of the combined module is equal to the accuracy of module that has been extracted directly from the original dataset.

#### Algorithm: Divide and Conquer algorithm

*Input:*

*S: A set of N number of sequences s1, s2, ……. sn*
*Threshold: Threshold for generating motifs*
*DistKey: distance key is threshold for creating a new cluster*
*Num-subs: Number of sub-sets in given data set*
*Output:*

*C: Cluster c1, c2 ……. ck*
*Algorithm:*

1. *generating M number of motifs m1, m2*, …….. *m M from given set of sequences using dynamic program*

~~~
  *with threshold and fixed length (L)*.
  *m1=substring of size L from fist sequence*
  *M=1 // number of motifs*
  *do until end of last sequence*
  *mi=next substring of size L*
        *for j=1 to no. of motifs*
        *find nearest motif exist*
        *end loop j*
        *if nearest motif exist then ith motif belongs to jth motif else*
        *M=M+1*
        *Create a new mth motif*
        *end if*
        *end do*
~~~
2. *generating motif frequency table using above motifs from given set of sequences*. ~~~
  *for i=1 to N*
  *for j=1 to M*
  *Freq-table(i,j) = number of times the jth motif appears in the ith sequences*
  *end loop j*
  *end loop i*
~~~
3. *dividing the given data into num_subs*
4. *for i=1 to num_subs*

~~~
  *Classifying the sub_sets into Ci classes using nearest-neighbor classifier with hamming distance measure and Dist_key*
  *end loop i*
~~~
5. *merging all classes generated in step 4* *Again classifying the all merged classes into final classes c1, c2* ……. *ck using nearest-neighbor classifies with hamming distance measure and Dist_key*

*Algorithm: Divide and Conquer classification of Grass genomes*

## 3. Experimental results

**Tabel-1:** [M-fold:10-times cross Validation: DAC experimental results-20000]

| GSD | DAC-Main Classification |  |  |  |  |  | DAC-Merge-Sub Classification |  |  |  |  |  |
| --- | --- | --- | --- | --- | --- | --- | --- | --- | --- | --- | --- | --- |
| DAC-validation | Total data | Training data | Training Time (S) | Testing Time(S) | Class | CA | Training data | Test data | Training Time(s) | Testing Time(s) | c-sub class | CA |
|  | 20000 | 12000 | 20 | 31 | 77 | 99.46 | 14 | 9 | 0.2 | 0.01 | 5 | 98.69 |
| Variance | 0 | 0 | 0.74 | .038 | 0.49 | 0.066 | 0.676 | 0.562 | 0.104 | 0.18 | 0.223 | 0.141 |

**Tabel-2:** [Medium size data: DAC experimental results-5000]

| GSD | DAC-Main Classification |  |  |  |  |  | DAC-Merge-Sub Classification |  |  |  |  |
| --- | --- | --- | --- | --- | --- | --- | --- | --- | --- | --- | --- |
|  | Total Data | Training Data | Training Time(S) | Testing Time(S) | Class | CA | Training data | Training Time(s) | Testing Time(S) | c-sub class | CA |
| GSD1 | 5000 | 3000 | 22 | 54 | 233 | 98.49 | 174 | 0.2 | 1 | 5 | 98.83 |
| GSD2 | 5000 | 3000 | 14 | 11 | 29 | 98.85 | 22 | 0.1 | 0.01 | 4 | 98.22 |
| GSD3 | 5000 | 3000 | 43 | 73 | 238 | 99.79 | 178 | 0.2 | 0.01 | 32 | 99.21 |
| GSD4 | 5000 | 3000 | 11 | 16 | 51 | 99.20 | 38 | 0.2 | 0.01 | 3 | 99.31 |
| GSD5 | 5000 | 3000 | 46 | 80 | 219 | 99.54 | 132 | 0.2 | 0.01 | 4 | 99.25 |

**Tabel-3:** [Large size data: DAC experimental results-20000]

| GSD | DAC-Main Classification |  |  |  |  |  | DAC-Merge-Sub Classification |  |  |  |  |
| --- | --- | --- | --- | --- | --- | --- | --- | --- | --- | --- | --- |
|  | Total data | Training Data | Training Time(S) | Testing Time(S) | Class | CA | Training data | Training Time(s) | Testing Time(S) | c-sub class | CA |
| DAC1 | 20000 | 12000 | 20 | 31 | 233 | 99.69 | 174 | 1 | 0.01 | 5 | 98.31 |
| DAC2 | 20000 | 12000 | 68 | 106 | 34 | 99.89 | 26 | 0.1 | 0.02 | 3 | 98.88 |
| DAC3 | 20000 | 12000 | 76 | 207 | 271 | 98.95 | 175 | 1 | 0.01 | 5 | 98.95 |
| DAC4 | 20000 | 12000 | 82 | 32 | 50 | 99.53 | 36 | 0.1 | 0.01 | 4 | 98.85 |
| DAC5 | 20000 | 12000 | 43 | 73 | 238 | 98.04 | 178 | 1 | 1 | 32 | 98.22 |

### 3.1. Data set: Large size data (100000)

There are 60000 patterns in the training set, 40000 patterns in the testing set and 826 classes. HDC is suitable for sequence data sets. To test the algorithms on large data sets, the number of training and test patterns were increased by updating one of the feature values in each pattern by a very small quantity. Results of sub classification reduced the training time as well as testing time, and increased classification accuracy.

### 3.2. Data set: Medium size data (25000)

There are 25000 patterns in the training set, 15000 patterns in the testing set and 154 classes. The results of Hamming Distance classifier (HDC), leader and Class—Subclass algorithms are as shown in Table 4 for medium data size. HDC is suitable for sequence data sets. To test the algorithms on Medium size data sets, the number of training and test patterns were increased by updating one of the feature values in each pattern by a very small quantity.

**Tabel-4:** [Large size data: Class With in the Class (CWC) - experimental results-20000]

| GSD | CWC-Main Classification |  |  |  |  |  | CWC-Sub Classification |  |  |  |  |
| --- | --- | --- | --- | --- | --- | --- | --- | --- | --- | --- | --- |
|  | Total Data | Training Data | Training Time(S) | Testing Time(S) | Class | CA | Training data | Training Time(s) | Testing Time(S) | c-sub class | CA |
| DAC1 | 20000 | 12000 | 17 | 12 | 14 | 99.95 | 9 | 0.2 | 0.01 | 4 | 94.83 |
| DAC2 | 20000 | 12000 | 19 | 36 | 65 | 99.46 | 45 | 0.2 | 0.01 | 18 | 94.74 |
| DAC3 | 20000 | 12000 | 14 | 15 | 16 | 99.93 | 11 | 0.2 | 0.01 | 9 | 88.81 |
| DAC4 | 20000 | 12000 | 19 | 15 | 16 | 99.91 | 11 | 0.2 | 0.01 | 9 | 88.85 |
| DAC5 | 20000 | 12000 | 19 | 36 | 65 | 99.46 | 45 | 0.2 | 0.01 | 18 | 94.74 |

### 3.3. Comparative experimental results

In order to validate the correctness of the methodology, a number of experiments were performed in comparison with other recorded methods. In the first case, the input dataset was derived from the 5 input set. The results are shown in Table 2. The number of splits was selected based on the size of the new dataset that would be produced each time, in order to maintain a similar processing time. It is obvious, that when the number of splits is n, the original dataset was processed. An improvement in the processing times can be seen from the table, while the accuracy is fairly constant. At this point, it must be noted that the number of the classes involved in each of the classification process is much larger due to the overlapping of the classes. The number of splits for each dataset is different to keep similar dataset sizes, in order to have comparable results. Results show a substantial improvement in the processing time while keeping almost constant model accuracy. The processing time in all cases follows the e-ax model, where a depends on the size of the original dataset and x is the number of splits, with minor fluctuations owing to the distribution of the instances of the overlapping grass genomes classes over the different dataset splits.

**Fig. 4.3.**
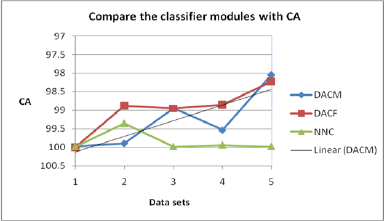
Comparison of CA of the Classifier Modules

**Fig. 4.4.**
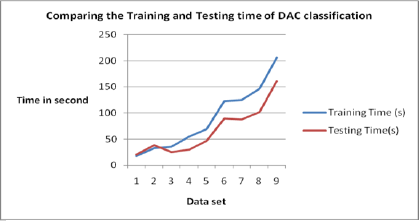
Comparison of Training and Test Time of DAC Classification

**Tabel-5:** Time complexity and classification accuracy of DAC for increasing size of data set:

| Data Set | DAC-Module Classification |  |  |  |  |  | DAC-Final Classification |  |  |  |  |
| --- | --- | --- | --- | --- | --- | --- | --- | --- | --- | --- | --- |
|  | TD | TrD | TrT(s) | TsT(s) | TnC | CA% | TrD | TrT(s) | TsT(s) | CsC | CA% |
| GSD 1 | 5000 | 3000 | 18 | 21 | 71 | 98.10 | 49 | 0.1 | 0.01 | 4 | 100 |
| GSD 2 | 10000 | 6000 | 123 | 90 | 91 | 99.99 | 60 | 0.1 | 0.01 | 14 | 96.66 |
| GSD 3 | 15000 | 9000 | 206 | 161 | 91 | 99.88 | 63 | 0.1 | 0.01 | 14 | 96.66 |
| GSD 4 | 20000 | 12000 | 33 | 39 | 22 | 99.97 | 15 | 0.1 | 0.1 | 6 | 100 |
| GSD 5 | 25000 | 15000 | 36 | 26 | 9 | 99.94 | 6 | 0.1 | 0.01 | 4 | 100 |
| GSD 6 | 30000 | 18000 | 55 | 30 | 9 | 99.53 | 6 | 0.1 | 0.01 | 4 | 100 |
| GSD 7 | 35000 | 21000 | 125 | 88 | 9 | 99.23 | 6 | 0.1 | 0.01 | 4 | 100 |
| GSD 8 | 40000 | 24000 | 146 | 102 | 9 | 99.33 | 6 | 0.1 | 0.01 | 4 | 100 |
| GSD 9 | 45000 | 27000 | 69 | 47 | 9 | 99.40 | 6 | 0.1 | 0.01 | 4 | 100 |

Note: 1. (GSD: Grass Genome Sequence Data Sets; TD: Total Data; TrD: Training data; TrT(s): Training Time(s); TsT(s): Testing Time(s); TnC: Total no of Class;CsC: C-sub class) 2. (Training data of DAC merged class is 2/3 of Total Number of classes)

### 3.4 Discussions

The analysis above shows that the running time and space requirement of our algorithm for computing *DAC* counts (for ranges of values of k) is dominated by the suffix array construction. This is especially true for the space requirement. To get an idea of whether our method can be applied to large sequence sets or not, we have to consider the space requirement of the suffix array constructions in more detail. The most space efficient suffix array construction requires (*n*⌈log2 *n*⌉)/8 bytes per input symbol plus 2*n*/8 bytes for representing the sequence. Given a 32-bit computer with 4 gigabytes (232 - 1 bytes) of main memory, *n* has to satisfy the inequality (2*n* + *n*⌈log2 *n*⌉)/8 ≤ 232 - 1. That is, the sequence length is limited to 1 gigabyte. Since we want to process considerably larger sequences, we developed a divide-and-conquer approach. This cuts the sequence into sufficiently small non-overlapping sections, such that for each section we can compute the corresponding enhanced suffix array on a 32-bit computer (equipped with 4 gigabytes of main memory). The representatives of the Subclass help in improving the CA and hence the Class–Subclass algorithm performs better than the leader algorithm. In bioinformatics (consisting of sequence data sets), it is required to find the subgroups/subfamilies in each of the grass genome group/family and Class-Subclass algorithm can be used for this application. The *Hamming Distance* problem has been described and the possible applications are mentioned. Though a part of the work was fund to have been done earlier (and therefore, rediscovered), the approach to obtain similarities as the inner product of the vectors representing the motifs enables one to use linear algebra techniques to reduce the cost of computation of similarities, and at the same time, keep the error as low as possible. By also taking into account the frequency of motifs in the pattern, the errors can be further reduced.

## 4. Conclusions

We have presented a novel approach for grass genome classification based on grid and parallel computing concept using Leader classifier and the hamming distance. Grass genome dataset is divided into multiple disjoint sets, where each one preserves the original class distribution. The new sets are then mined in parallel for knowledge, using leader classification algorithm, and the extracted knowledge modules are combined in final classes. Results indicate that the proposed method is time efficient and shows that overall accuracy is comparable with other methods. It must be noted that the parallelization of the procedure allow the processing of much larger datasets as compared to other techniques. The representatives of the subclasses help in improving the CA. The Class-Subclass algorithm performs better than the leader algorithm.

